# Mitochondrial Proteome of Affected Neurons in a Mouse Model of Leigh Syndrome

**DOI:** 10.1101/2019.12.29.890541

**Authors:** Alejandro Gella, Patricia Prada-Dacasa, Montserrat Carrascal, Melania González-Torres, Joaquin Abian, Elisenda Sanz, Albert Quintana

## Abstract

Defects in mitochondrial function lead to severe neuromuscular orphan pathologies known as mitochondrial disease. Among them, Leigh Syndrome is the most common pediatric presentation, characterized by symmetrical brain lesions, hypotonia, motor and respiratory deficits, and premature death. Mitochondrial diseases are characterized by a marked anatomical and cellular specificity. However, the molecular determinants for this susceptibility are currently unknown, hindering the efforts to find an effective treatment. Due to the complex crosstalk between mitochondria and their supporting cell, strategies to assess the underlying alterations in affected cell types in the context of mitochondrial dysfunction are critical. Here, we developed a novel virus-based tool, the AAV-mitoTag viral vector, to isolate mitochondria from genetically-defined cell types. Administration of the AAV-mitoTag in the vestibular neurons of a mouse model of Leigh Syndrome lacking the complex I subunit *Ndufs4* allowed us to assess the proteome and acetylome of susceptible neurons in a well characterized model recapitulating the human disease. Our results show a marked reduction of complex-I N-module subunit abundance and an increase in the levels of the assembly factor NDUFA2. Transiently-associated non-mitochondrial proteins such as PKCδ, and the complement subcomponent C1Q were also increased in *Ndufs4-deficient* mitochondria. Furthermore, lack of *Ndufs4* induced pyruvate dehydrogenase (PDH) subunit hyperacetylation, leading to decreased PDH activity. We provide novel insight on the pathways involved in mitochondrial disease, which could underlie potential therapeutic approaches for these pathologies.

## 1 Introduction

Severe alterations of the mitochondrial machinery involved in energy generation lead to a group of progressive and usually fatal pathologies collectively known as primary mitochondrial disease (MD), affecting 1:5000 births (Schaefer et al., 2004). Currently, there is no cure and the treatments available are mostly ineffective (Schon et al., 2010). MD predominantly affects high energy-requiring organs such as the brain (Vafai and Mootha, 2012). Indeed, neurological damage plays a prominent role in MD pathology and lethality (DiMauro and Schon, 2008). However, not all neuronal populations are equally vulnerable to MD, but rather show a striking anatomical and cellular specificity (Dubinsky, 2009). Recent reports have evidenced the remarkable regional heterogeneity in mitochondrial protein composition in the brain, which may account for the cell type specificity of MD (Pagliarini et al., 2008)

Humans harboring mutations in NDUFS4, a subunit of the mitochondrial complex I involved in the assembly and stability of the complex (Scacco et al., 2003; Kruse et al., 2008; Calvaruso et al., 2012) develop Leigh Syndrome (LS), a severe form of MD (DiMauro and Schon, 2008). These individuals die at an early age, commonly presenting brain lesions, hypotonia and respiratory deficits (Leshinsky-Silver et al., 2009; Lombardo et al., 2014). Mice lacking this protein (NDUFS4KO) recapitulate the human symptoms and are an excellent correlate of the human pathology (Quintana et al., 2010; Breuer et al., 2013). As observed in LS patients, NDUFS4 deficiency in mice leads to prominent bilateral lesions in the dorsal brainstem, more specifically in the vestibular nucleus (VN). Conditional NDUFS4 ablation restricted to the VN causes a pathology similar to global Ndufs4KO mice. Conversely, viral-mediated re-expression of *Ndufs4* in the VN is sufficient to ameliorate the pathology and to extend the lifespan of Ndufs4KO mice (Quintana et al., 2012a). In addition, NDUFS4 deficiency reveals a cell type-specific susceptibility to the disease. NDUFS4 ablation in glutamatergic neurons (Vglut2-expressing) recapitulates most of the phenotype present in the global NDUFS4-deficient mice (Bolea et al., 2019), thus warranting further study of glutamatergic neurons in the VN.

Here, we have developed and implemented a novel viral vector approach (AAV-mitoTag) to purify cell type-specific mitochondria in a fast and minimally-disruptive manner from glutamatergic neurons in the VN. This approach has allowed us to gain a better understanding of both mitochondrial proteome and acetylome specifically in cells highly susceptible to mitochondrial dysfunction. Hence, we provide insight on the specific proteomic changes driven by NDUFS4 deficiency, which may underlie the molecular and metabolic alterations elicited by mitochondrial dysfunction, with the overarching goal of identifying novel molecular targets with therapeutic potential for LS and MD.

## 2 Material & Methods

### Animal husbandry

Mice were maintained with Teklad Global rodent diet No. 2018S (HSD Teklad Inc., Madison, Wis.) and water available *ad libitum* in a vivarium with a 12-hour light/dark cycle at 22°C. All experiments were conducted following the recommendations in the Guide for the Care and Use of Laboratory Animals and were approved by the Animal Care and Use Committee of the Universitat Autònoma de Barcelona (Barcelona, Spain).

### Mouse genetics

*Slc17a6^Cre^* (BAC-Vglut2::Cre) mice were generated by Ole Kiehn (Borgius et al., 2010). *Ndufs4*^lox/lox^ and *Ndufs4*^Δ/+^ were previously generated by our group (Kruse et al., 2008; Quintana et al., 2010). Mice with conditional deletion of *Ndufs4* in Vglut2-expressing glutamatergic neurons (*Slc17a6*^Cre^, *Ndufs4*^Δ/lox^ or Vglut2:Ndufs4cKO) were generated by crossing mice with one *Ndufs4* allele deleted and expressing a codon-improved Cre recombinase (iCre) under the *Slc17a6* promoter (*Slc17a6*^Cre^, *Ndufs4*^Δ/+^) to mice with two floxed *Ndufs4* alleles (*Ndufs4*^lox/lox^). Littermate controls were *Slc17a6*^Cre^, *Ndufs4*^lox/+^ (Vglut2:Ndufs4cCT).

### Cell cultures

HEK293T cells were obtained from the American Type Culture Collection (ATCC). Cells were grown in high glucose Dulbecco’s Modified Eagle Medium (Gibco) supplemented with 10% (v/v) fetal calf serum (Gibco), l-glutamine (Gibco) and penicillin/streptomycin (Gibco) and maintained at 37 °C with 5% CO_2_. Cells were seeded at 200,000 cells/mL and transfected with plasmids with calcium phosphate and/or transduced with adeno-associated viral vectors (AAVs) for 4 days prior to immunostaining for hemagglutinin (HA), western blot analysis or mitochondria immunoisolation (mitoTag) assays.

### Immunofluorescence

For immunofluorescence analysis of cultured cells, HEK293T cells were fixed in 4% paraformaldehyde (PFA) for 10 min at RT, and further washed two times with PBS before permeabilization with 0.25% Triton X-100 in PBS for 10 min. After permeabilization, cells were washed three times for 5 min with PBS, blocked with 1% BSA in PBS-Tween 20 (0.05%) for 60 min, and incubated with anti-HA antibodies (at 1:1000 dilution, #901514 BioLegend) overnight at 4 °C. After incubation, cells were washed three times with PBS for 5 min and further incubated with anti-mouse Alexa 555 antibody (1:500, Thermo Fisher Scientific). Finally, cells were washed three times with PBS for 5 min, counterstained with Hoechst 33258 (2 μg/ml, Sigma-Aldrich) in PBS for 10 min and mounted onto slides using Fluoromount G (Electron Microscopy Sciences) before microscopic analysis. Immunofluorescence analysis was accomplished in mouse brains fixed overnight in 4% PFA in PBS (pH 7.4). Subsequently, brains were cryoprotected in a PBS solution containing 30% (w/v) sucrose and frozen in dry ice. Frozen brains were embedded in OCT, sectioned at 30 μm in a cryostat and rinsed in PBS-Tx buffer (Phosphate-buffered saline containing 0.2% (v/v) Triton X-100). Non-specific binding was blocked with 10% (v/v) normal donkey serum (NDS) in a PBS-Tx solution for 60 min at room temperature, followed by overnight incubation at 4°C with primary antibodies diluted in 1% NDS-PBS-Tx (1:1,000 for mouse anti-HA, #901514 BioLegend; 1:1,000 for rabbit anti-TOM20, #sc-11415 Santa Cruz Biotech). Sections were then washed in PBST and incubated for 1 h at room temperature with the corresponding Alexa Fluor-conjugated secondary antibodies (1:500, Thermo Fisher Scientific) in 1% NDS-PBS-Tx. Sections were finally washed in PBS-Tx and rinsed in water before mounting onto slides with Fluoromount G with DAPI (Electron Microscopy Sciences) for microscopic analysis.

### Western Blotting

For western blot analysis of cultured cells, HEK293T cells were lysed with SDS sample buffer (62.5 mM Tris–HCl, pH 6.8, 2% SDS, 10% glycerol, all from Sigma-Aldrich) without dithiothreitol (DTT) and bromophenol blue, to avoid interference with protein quantification. Prior to protein determination, samples were sonicated to shear genomic DNA and reduce the viscosity of the lysates. Protein concentration of the lysates was determined using the BCA protein assay (Thermo Fisher Sci.) according to the manufacturer’s instructions. After protein quantification, 50 mM DTT and 0.1% bromophenol blue (Sigma-Aldrich) were added to the samples. Western blot analysis of whole tissue lysates or immunoprecipitates from mitoTag assays was accomplished by adding 4X laemmli sample buffer to samples prior to heat-denaturing. Protein lysates (10 μg for HEK293T cell lysates; 0.125% whole tissue lysates and 10% immunoprecipitates for mitoTag assays) were heated at 99 °C for 3 minutes and subjected to 4-20% gradient SDS-PAGE prior to transfer to nitrocellulose membranes (Bio-Rad Laboratories, Inc.). Membranes were then blocked for 1 hour with 5% (w/v) dried skimmed milk in TBS-T buffer (Tris-buffered saline containing 0.1% (v/v) Tween-20) and incubated overnight at 4°C with primary antibodies against VDAC1 (1:1,000, #ab18988 Abcam), Calreticulin (1:1,000, #ab2707 Abcam), TOM20 (1:5,000, #sc11415 Santa Cruz Biotech), β-actin (1:40,000, #A1978 Sigma). After incubation with the corresponding HRP-conjugated secondary antibodies (1:10,000; Jackson ImmunoResearch), membranes were washed in TBS-T and developed using an enhanced chemiluminescence (ECL) detection system (Thermo Fisher Sci.). Bands were quantified using Image J software (National Institutes of Health).

### Transmission Electron Microscopy

Vestibular nuclei mitochondria were fixed in 2.5% (v/v) glutaraldehyde (Merck) in 0.1 M phosphate buffer (PB), pH 7.4 at 4°C overnight. Postfixed for 2 h with 1% (w/v) osmium tetroxide (TAAB Lab) containing 0.8% (w/v) potassium hexacyanoferrate (Sigma-Aldrich) in PB, dehydrated in gradual ethanol (50-100%), embedded in Eponate 12 resin (Ted Pella Inc.), and polymerized at 60°C for 48 h. Ultrathin sections of 50 nm were collected onto carbon-coated copper grids of 200 mesh and contrasted with conventional uranyl acetate and lead citrate solutions. Sections were visualized in a TEM Jeol JEM-1400 (Jeol Ltd., Tokyo, Japan) operating at 80 kV and equipped with a CCD Gatan ES1000W Erlangshen camera (Gatan Inc.).

### MitoTag Adeno-Associated viral vector (AAV) generation

We generated an adeno-associated virus (AAV) that would initiate Cre-inducible mitoTag expression using a double floxed inverted open reading frame (DIO) approach. An NheI/AscI fragment containing a construct coding for the mouse *Tomm20* gene fused to three hemagglutinin (HA) sequences (Tomm20·3HA) was cloned into a NheI/AscI-digested pAAV-EF1a-DIO-WPRE-hGH polyA plasmid to obtain a pAAV1-EF1a-DIO-Tomm20·3HA-WPRE-hGH polyA construct (AAV-mitoTag). An AAV (AAV1 serotype) vector was produced and CsCl-gradient purified as described previously (Quintana et al., 2012b). The TOM20-HA fusion protein was generated by mutation of *Tomm20* stop codon to glutamic acid (E) and phenylalanine (F).

### AAV-mitoTag delivery

Mice (n=28, 37.4±7 days, 22.1±3.2 g) were anaesthetized with isoflurane (5.0 % induction, 1.5-0.7 % maintenance) and placed in a stereotaxic frame (Kopf Instruments). Ketoprofen analgesic solution (5 mg/kg s.c; Sanofi-aventis) and ocular protective gel (Viscotears^®^) were applied before stereotaxic procedure. Mice received intracranial AAV injections into the vestibular nucleus (VN) at the following coordinates: AP −0.6, ML ± 0.125, DV −0.39 mm from the Bregma, using the correction factor (Bregma - Lambda/4.21). A total of 1.0 μL of AAV-mitoTag vector (0.7 × 10^12^ viral genomes/mL; 0.5 μL per side) was injected into each hemisphere at a rate of 0.25 μL/min using a 5 μL Hamilton syringe with a 32G blunt needle. Following injection, the needle was kept in place for 4 minutes post-delivery to allow proper viral vector diffusion and, subsequently, removed at 1 mm/min.

***Isolation of mitochondria by immunoaffinity purification***

For HEK293T cells, 200,000 cells/mL were transduced with AAV-mitoTag and a viral vector expressing Cre recombinase (AAV-Pkg-Cre; UNC Vector Core) or left untransduced for 4 days prior to lysis in mitochondria isolation kit lysis buffer (1×10^7^ cells/mL; Miltenyi Biotec) with protease inhibitors (Sigma-Aldrich) and homogenized by passing the lysate 25 times through a 1 mL syringe with a 25G needle. Homogenized lysates were next centrifuged at 700 x g for 10 min at 4°C, and supernatants collected and incubated with anti-HA or anti-TOM22 microbeads (Miltenyi Biotec) according to manufacturer’s directions. Mitochondria from glutamatergic cells in the vestibular nucleus (VN) were isolated from freshly collected mouse brain tissue. Tissue homogenates were prepared using the Mitochondria Extraction Kit, tissue (Miltenyi Biotec) according to manufacturer’s instructions. Briefly, following decapitation, brains were weighted, and vestibular nuclei dissected according to the Paxinos mouse brain atlas and digested with extraction buffer for 30 min at 4°C. After centrifugation at 300 × g for 5 min at 4°C, pellets were resuspended in Solution 2 supplemented with EDTA-free protease inhibitor cocktail (Roche Diagnostics), and deacetylase inhibitors (10 mM nicotinamide, 10 μM TSA and 10 mM sodium butyrate). Digested tissue was homogenized using a glass dounce homogenizer. Homogenates were centrifuged at 1,000 × g for 5 min at 4°C. Subsequently, supernatants containing mitochondria were further purified as directed in the Mitochondria Isolation Kit (Miltenyi Biotec). For magnetic labeling and isolation, supernatants diluted in Separation Buffer were incubated with 50 μL of anti-HA Microbeads on a shaker (60 rpm) for 1 h at 4 °C. Subsequently, LS columns (Miltenyi Biotec) were placed in a magnetic QuadroMACS™ Separator (Miltenyi Biotec) and equilibrated with 3 mL Separation Buffer (Miltenyi Biotec). Microbead-coated mitochondria were applied onto the LS column, followed by three 3-mL washing steps with Separation Buffer. Columns were removed from the separator and mitochondria gently flushed out in 1.5 mL of Separation Buffer with a plunger. Mitochondria were pelleted by centrifugation for 2 minutes at 12,000 x g and washed twice with Storage Buffer. Two immunoprecipitations (from 4-5 mice) were pooled for proteomic analysis.

***Isolation of mitochondria by double centrifugation***

Mitochondria were isolated by differential centrifugation using protocols previously described (Brown et al., 2004; Pagliarini et al., 2008). Briefly, following decapitation, vestibular nuclei were dissected according to the Paxinos mouse brain atlas and homogenized using a glass dounce homogenizer containing MAS buffer (70 mM sucrose, 220 mM mannitol, 10 mM KH2PO4, 5 mM MgCl_2_, 1 mM EGTA, 2 mM HEPES-KOH, pH 7.2). Homogenates were centrifuged at 900 × g for 10 min at 4°C. Supernatants were further centrifuged at 9,000 × g for 10 min at 4°C. Resulting supernatants were discarded and pellets containing mitochondria were resuspended in MAS buffer. Protein concentration was determined using the BCA protein assay (Thermo Fisher Sci.) according to the manufacturer’s instructions.

### PDH enzyme activity

PDH activity was measured in VN homogenates by using the Pyruvate Dehydrogenase (PDH) Enzyme Activity Microplate Assay Kit (Abcam) according to manufacturer’s instructions. Briefly, mitochondria isolated by double centrifugation (50 μg) were loaded into the wells of a 96-well plate that was specific for PDH activity assay. PDH enzyme was immunocaptured by the monoclonal antibodies coated on the wall of each well. The enzymatic activity of PDH was determined based on the production of NADH coupled to a reporter dye whose formation is monitored by measuring the increase in absorbance at 450 nm on a Varioskan LUX plate reader (Thermo Fisher Sci.).

### Measurement of respiration in isolated mitochondria

Oxygen consumption rate (OCR) was measured in MAS buffer supplemented with 0.5 % (w/v) fatty acid–free BSA, using a Seahorse XFp Extracellular Flux Analyzer (Agilent Technologies, Inc). Freshly isolated mitochondria (5-20 μg of protein) obtained by double centrifugation were plated in each well of an XFp plate in 25 μL of MAS-BSA Buffer. To attach mitochondria, the plate was centrifuged at 2,000 x g at 4°C for 20 min and then 155 μL of MAS-BSA buffer containing 10 mM pyruvate/5 mM malate, 100 μM palmitoyl-L carnitine/5 mM malate, or 20 mM succinate/4 μM rotenone was added to each well. First, basal measurements of oxygen consumption rate (state 2) were obtained. Next, ADP (5 mM) was injected (state 3), followed by sequential injections of oligomycin (2 μM, state 4o), FCCP (5 μM, state 3u) and antimycin A/rotenone (2.5 μM and 2 μM respectively) to disrupt mitochondrial respiration. Together, this injection series allowed for the determination of ADP-uncoupled respiration (ADP), proton leak (oligomycin), maximal respiration (FCCP) and non-mitochondrial residual oxygen consumption (antimycin A/rotenone).

### Protein extraction and digestion

Mitochondria from glutamatergic neurons in the vestibular nucleus were immunocaptured as described above. These mitochondria were then suspended in lysis buffer (4% (w/v) SDS, 100 mM Tris-HCl, pH 7.6, 0.1 M DTT) and incubated at 95°C for 5 min. Protein extracts were quantified using RC-DC^TM^ Protein Assay (Bio-Rad). Three biological replicates from Vglut2:Ndufs4cCT and Vglut2:Ndufs4cKO mice mitochondria were processed. Protein was digested with sequencing grade modified trypsin (Promega) using the FASP (Filter Aided Sample Preparation) digestion protocol(Wisniewski et al., 2009). Briefly, 150 μg of protein from each sample were loaded onto a 10-kDa Amicon Ultra-0.5 centrifugal filter (Millipore). The protein mixture was washed three times by adding 200 μL of UA buffer (8 M Urea, 0.1 M Tris-HCl pH 8.5) to the filter and centrifuging at 14,000×g for 10 min at 13°C. Next, proteins were alkylated with 100 μL of alkylation buffer (0.05 M IAA, 8 M Urea, 0.1 M Tris-HCl pH 8.5) in the dark for 20 min at 25°C. Subsequently, the protein extract was washed three times with 100 μL of UA buffer and three times with 100 μL of 200 mM triethylammonium bicarbonate (TEAB). Trypsin digestion was performed at 37°C for 18 h using an enzyme-to-protein ratio of 5:100. Tryptic peptides were eluted by the addition of 3 × 100 μL of 200 mM TEAB followed by a centrifugation at 14,000 × g for 15 min at 13 °C.

### Peptide labeling by isobaric tandem mass tag

Each tryptic peptide mixture obtained from 150 μg of protein extract was isotopically labeled with the corresponding TMT6plex reagent (Thermo Fisher Sci.) based on the standard procedure. TMT-labeled peptide mixtures were combined in a low-bind 1.5 mL Eppendorf tube, evaporated, and desalted using a C18 SPE cartridge (Agilent Technologies). The SPE eluates were evaporated and resuspended in 50 mM MOPS pH 7.2, 10 mM Na2HPO4, 50 mM NaCl.

### Acetylated Peptide Enrichment

The TMT-labeled peptide mixtures (900 μg) were enriched for acetylated peptides by immunoaffinity purification using an anti-acetyl lysine antibody following the PTMScan^®^ Pilot Acetyl-Lysine Motif (Ac-K) protocol (Cell Signaling Technologies). Peptides eluted from the enrichment were desalted using PolyLC C18 tips (PolyLC Inc., MD, USA), and resuspended in 5% methanol (1% formic acid) for mass spectrometry analysis.

### LC—MSn Analysis

Peptides were analyzed using an Orbitrap Fusion Lumos Tribrid mass spectrometer (Thermo Fisher Sci.) equipped with a Thermo Scientific Dionex Ultimate 3000 ultrahigh-pressure chromatographic system (Thermo Fisher Sci.) and Advion TriVersa NanoMate (Advion Biosciences Inc.) as the nanospray interface. Peptide mixtures were loaded to a μ-precolumn Acclaim C18 PepMap100 (100 μm × 2 cm, 5 μm, 100 Å, C18 Trap column; Thermo Fisher Sci.) at a flow rate of 15 μL/min and separated using a C18 Acclaim PepMap RSLC analytical column (75 μm × 50 cm, 2 μm, 100 Å, nanoViper). Separations were done at 200 nL/min using a linear ACN gradient from 0 to 40% B in 240 min (solvent A = 0.1% formic acid in water, solvent B = 0.1% formic acid in acetonitrile). The mass spectrometer was set up in the positive ion mode and the analysis was performed in an automatic data dependent mode (a full scan followed of 10 HCD scan for the most abundant signals).

### Database search and proteomic data analysis

Database search was done using Proteome Discoverer v2.1 (Thermo-Instruments) with a 1% false discovery rate (FDR) and the UniProt 2018-10 database restricted to *Mus musculus* (mouse) and contaminants. Search parameters were precursor and fragment tolerance, 20 ppm; enzyme, trypsin; missed cleavages, 1; fixed modifications, TMTsixplex (N-terminal, K), carbamidomethyl (C); and variable modifications, oxidation (M).

For proteomic data analysis, DanteR (Taverner et al., 2012) was used for relative quantification. Only unique peptides were considered for the analysis. Tandem mass tag (TMT) reporter intensity data were normalized using the QuantileN method and ANOVA analysis was performed. In the acetylome quantification, the ratio of the quantified peptides was normalized using the ratio of the corresponding protein. Differential proteins and peptides were selected using a p-value cutoff of 0.05 and a fold change lower than 0.7 (down) or higher than 1.3 (up).

The mass spectrometry proteomic data have been deposited to the ProteomeXchange Consortium (http://proteomecentral.proteomexchange.org) via the PRIDE (Vizcaino et al., 2016) partner repository with the dataset identifier PXD016321.

### Bioinformatics Analysis

Functional enrichment analysis using overrepresentation analysis (ORA) with the Gene Ontology (Biological process) and Pathway (Reactome) databases was accomplished using WebGestalt (Liao et al., 2019) (http://webgestalt.org/). Affinity propagation was used for redundancy reduction. The STRING (Search Tool for the Retrieval of Interacting Genes/Proteins, https://string-db.org/) algorithm (version 11) was used to build protein–protein interaction networks. Only high-confidence interactions (score 0.7) were chosen.

### Statistics

Data are shown as the mean ± SEM. GraphPad Prism v5.0 software was used for statistical analyses. Appropriate tests were selected depending on the experimental design as stated in the figure legends. Statistical significance, when reached *(P* <0.05 was considered significant), is stated in the figure legends.

## 3 Results

### Generation and validation of the AAV-mitoTag viral vector

To allow the isolation of the mitochondria from susceptible neuronal cell types, a Cre-dependent mitoTag adenoassociated viral vector (AAV-mitoTag) was generated by inserting the cDNA of an HA-tagged *Tomm20* into a double-floxed inverse open reading frame (DIO) AAV construct (Fig. 1A). Analysis for Cre recombinase-dependency of the construct (Fig. 1B) and mitochondrial targeting of the resulting TOM20·HA fusion protein (Fig. 1C) was first assessed in HEK293T/17 cells. Transfection of the viral vector construct showed that the TOM20·HA fusion protein was only expressed in cells transduced with a viral vector expressing Cre recombinase. In these cells, fusion protein showed the expected molecular weight. Immunofluorescence analysis for HA in cells transfected with the mitoTag viral vector construct and transduced with viral vector particles expressing Cre recombinase showed a localization of the fusion protein consistent with incorporation into mitochondria (Fig. 1C). To test whether whole mitochondria isolation from Cre-expressing cells could be accomplished by an immunocapture method, lysates from HEK293T/17 cells transduced with the AAV-mitoTag viral vector (AAV-DIO-Tomm20·HA) along with a Cre recombinase-expressing viral vector (AAV-Cre) were used to immunoprecipitate HA-tagged mitochondria with anti-HA coated magnetic microbeads (Fig. 1D). Anti-TOM22 coated magnetic microbeads were also used as control to immunoprecipitate unlabeled mitochondria from non-transduced cells. Western blot analysis of the immunoprecipitates showed specific immunoisolation of mitochondria without contamination from cytosolic proteins.

**Figure 1.**
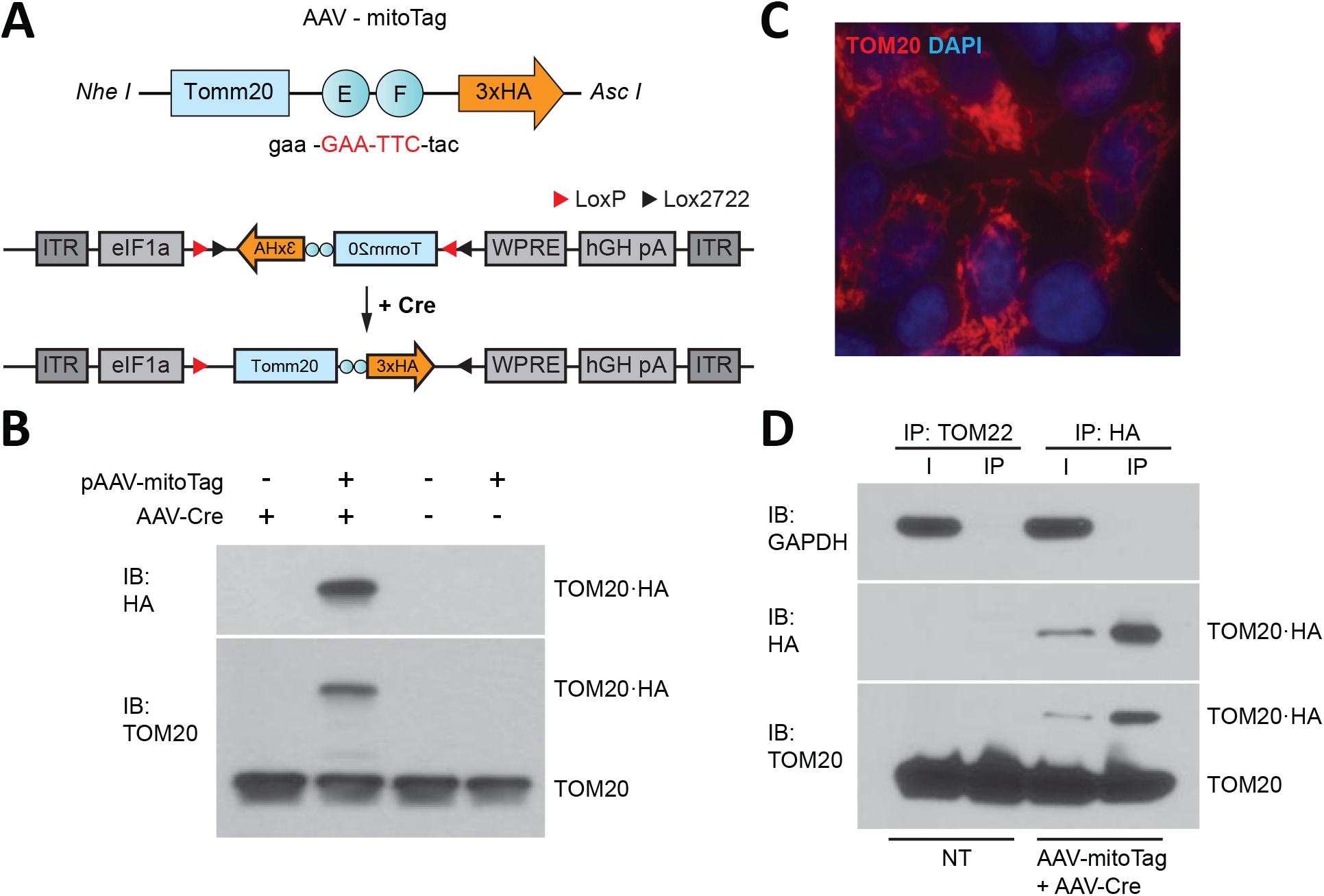
Generation of the mitoTag viral vector (AAV-mitoTag). (A) A TOM20·hemagglutinin (HA) fusion protein was generated by mutation of *Tomm2O* stop codon (red). The resulting construct was cloned in a Cre-dependent viral vector plasmid (pAAV-DIO). (B) Western blot analysis showing the expression and Cre-dependency of the pAAV-DIO-TOM20·HA construct in HEK293T cells. Immunoblot (IB) for HA and TOM20 shows expression of the fusion protein TOM20·HA at the expected molecular weight only in cells transduced with the Cre-expressing viral vector (AAV-Cre). (C) Immunofluorescence analysis showing mitochondrial localization of TOM20·HA in HEK293T cells. Cells were transfected with the pAAV-DIO-TOM20·HA construct and transduced with a viral vector expressing Cre recombinase (AAV-Cre) prior to fixation and immunostaining using an antiHA antibody. (D) Western blot analysis showing specific immunoprecipitation of mitochondria using anti-HA magnetic microbeads. GAPDH was used as a cytosolic marker. TOM20 was used as a mitochondrial marker. Immunoprecipitation assays with anti-TOM22 microbeads were used as a control for mitochondria isolation in non-transduced cells.

### Cell type-specific isolation of mitochondria from glutamatergic neurons in the VN

Glutamatergic neurons (Vglut2-expressing) in the VN have been proven susceptible to complex I deficiency (Bolea et al., 2019). To define the factors driving this specific susceptibility, we first confirmed the feasibility of the AAV-mitoTag approach for the isolation of mitochondria from this genetically defined neuronal population *in vivo.* To show expression of the mitoTag in mitochondria from the cell type of interest, AAV-mitoTag viral vector was injected bilaterally into the VN of *Slc17a6*^Cre^ (Vglut2^Cre^) mice. Subsequently, brain sections containing the area were analyzed by immunofluorescence using anti-HA antibodies (Fig. 2A). HA staining was restricted to specific cells in the VN, and double-labeling analysis using anti-HA and anti-TOM20 antibodies confirmed the mitochondrial localization of the mitoTag (Fig. 2B). Immunocapture of mitochondria from Vglut2-expressing cells was accomplished from lysates of pooled vestibular nuclei using Anti-HA coated magnetic microbeads (Fig. 2C). Western blot analysis of whole lysates (input of the immunoprecipitation) and immunoprecipitates showed specific enrichment for mitochondrial markers such as the mitochondrial porin VDAC1 or the translocase of the outer mitochondrial membrane TOM20. In contrast, cytosolic markers such as the cytoskeleton protein beta-Actin or endoplamatic reticulum (ER) markers such as calreticulin, appeared depleted in the IP, confirming mitochondrial enrichment *in vivo* with the mitoTag approach (Fig. 2D). Electron microscopy (EM) analysis of the immunoprecipitates from VN lysates of Vglut2^Cre^ mice injected with the mitoTag viral vector confirmed binding of the anti-HA magnetic microbeads to the outer membrane of whole mitochondria (Fig. 2E).

**Figure 2.**
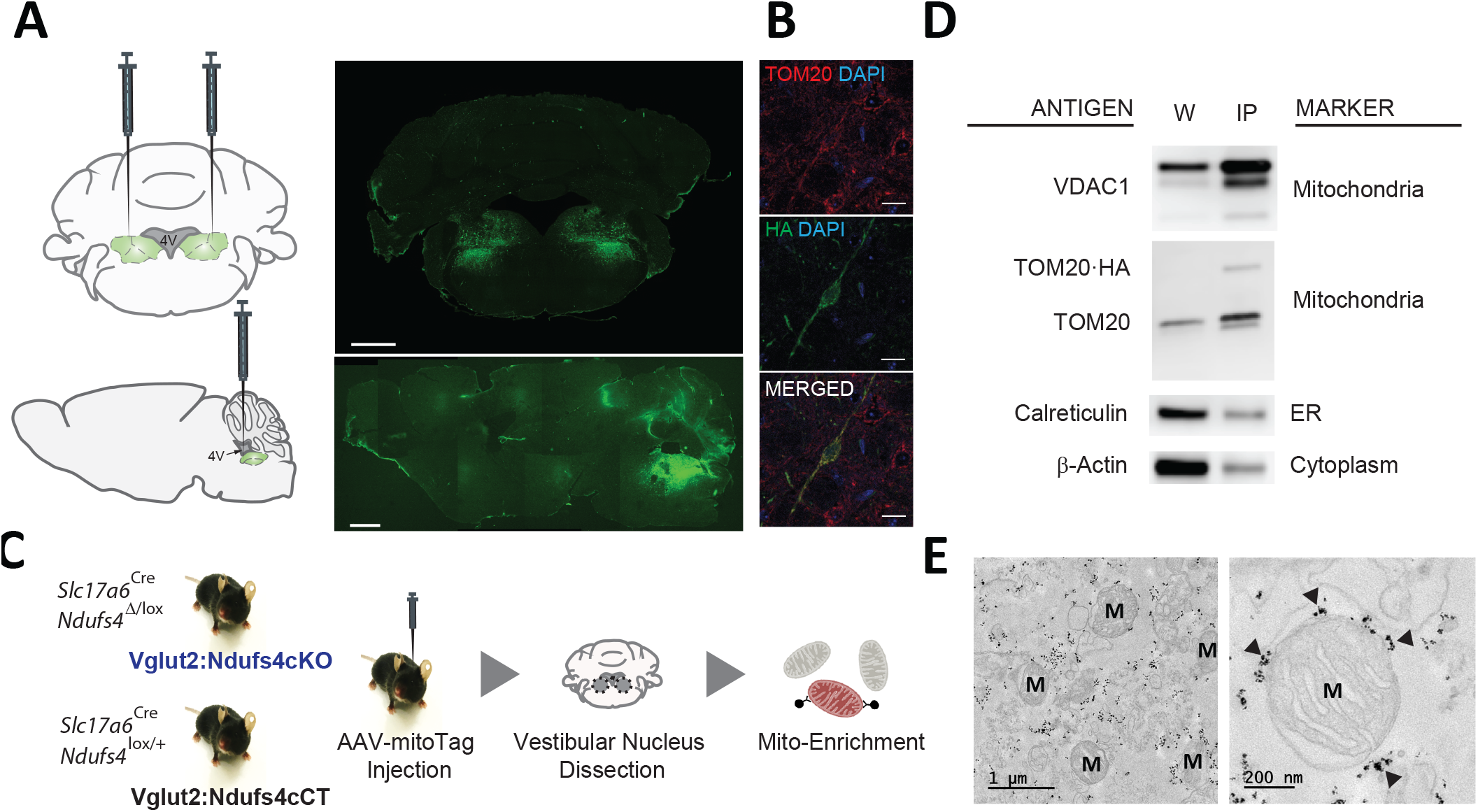
*In vivo* validation of the mitoTag approach. (A) Diagram of the targeted injection site (VN; coronal and sagittal view) and immunofluorescence analysis using an anti-HA antibody to confirm transduced neurons in the VN of Vglut2^Cre^ mice. Scale bar: 1 mm (B) Double immunostaining for TOM20 and TOM20·HA showing mitochondrial localization of the fusion protein. Scale bar: 50 μm (C) Cartoon showing the experimental approach. (E) Western blot analysis for organelle-specific proteins in whole-cell lysates (W) and lysates of mitoTag immunoprecipitates (IP). ER, endoplasmic reticulum (F) Electron micrographs of mitoTag preparations from the VN of Vglut2:Ndufs4cCT mice. Electron dense spots (black arrows) are anti-HA magnetic microbeads bound to the outer mitochondrial membrane. M, mitochondria.

### Identification and quantitation of the global and acetyl-proteome in mitochondria from glutamatergic neurons in the VN

Next, we sought to define the specific protein changes induced by complex I deficiency in mitochondria from glutamatergic neurons in the VN. To accomplish this, mice conditionally deficient for the complex I subunit NDUFS4 in glutamatergic neurons (Vglut2:Ndusf4cKO mice) and their controls (Vglut2:Ndusf4cCT mice) were injected with the AAV-mitoTag viral vector to isolate mitochondria from glutamatergic neurons in the VN (Fig. 2C). Cell type-specific analysis of the global mitochondrial proteome and acetylproteome from Vglut2:Ndufs4cKO and Vglut2:Ndufs4cCT mice was accomplished by liquid chromatography-tandem mass spectrometry (LC–MSn). Global and acetylome were simultaneously identified and quantified from pooled mitoTag preparations from the VN of Vglut2:Ndufs4cKO (n=3) and Vglut2:Ndufs4cCT mice (n=3). Each replicate was a pool of two mitoTag preparations. 95% of labelled peptides were dedicated to analysis of the acetylproteome, the remaining 5% was used for global proteome analysis.

For global proteome analysis, a total of 34,297 spectra corresponding to 17,636 non-redundant peptides were identified through database search (1% FDR). Only peptides identified as unique (i.e., peptide sequences belonging to one single protein in the database) were considered. Overall, a total of 3,037 proteins were quantified from 16,042 non-redundant unique peptides. Among these, 40 proteins showed increased abundance (> 1.2-fold, p-value ≤ 0.05), while 25 proteins presented reduced levels (< 0.8-fold, p-value ≤ 0.05) in mitoTag preparations from Vglut2:Ndufs4cKO mice when compared to controls. Remarkably, most complex I subunits (34 out of 45 detected proteins) showed significantly reduced levels in NDUFS4-deficient mitochondria, except for the assembly factor NDUFAF2, which showed increased abundance in these mitochondrial preparations (Fig. 3A-B). Constituents of the NADH dehydrogenase module (N-module: NDUFA2, NDUFA12, NDUFV1, NDUFV2, NDUFV3, NDUFS1, NDUFS6 and NDUFS4) were the most significantly decreased proteins in these mitochondria (Fig. 3B). Constituents of the Q-module, which is located adjacent to N-module, and specific subunits of the P-Module, were also reduced in Vglut2:Ndufs4CKO VN mitochondria (Fig. 3B). Proteins with increased abundance in these mitochondria included the A, B and C-chain polypeptides of the complement subcomponent C1q (C1QA, C1QB, C1QC), which are known to bind to the mitochondrial protein gC1qR (globular head C1q receptor or C1QBP) to modulate mitochondrial metabolism (Ling et al., 2018), and protein kinase C delta (PKCδ), a member of the protein kinase C family of serine- and threonine-specific protein kinases that translocates to mitochondria upon different stimuli (Majumder et al., 2001). The beta and gamma1 subunits of the glycogenolytic phosphorylase kinase complex (PHKB and PHKG1) also appeared increased. Functional enrichment analysis for overrepresented biological processes (Fig. 3C) and pathways (Fig. 3D) confirmed complement-mediated signaling and glycogen metabolism as the most enriched protein sets (terms) in the analysis of the proteins with increased abundance in mitochondria from the VN of Vglut2:Ndufs4cKO mice. Analysis of underrepresented proteins resulted in highly significant categories confirming a role in mitochondrial organization, and more specifically in respiratory electron transport and complex I biogenesis for the proteins that showed reduced presence in NDUFS4-deficient mitochondria.

**Figure 3.**
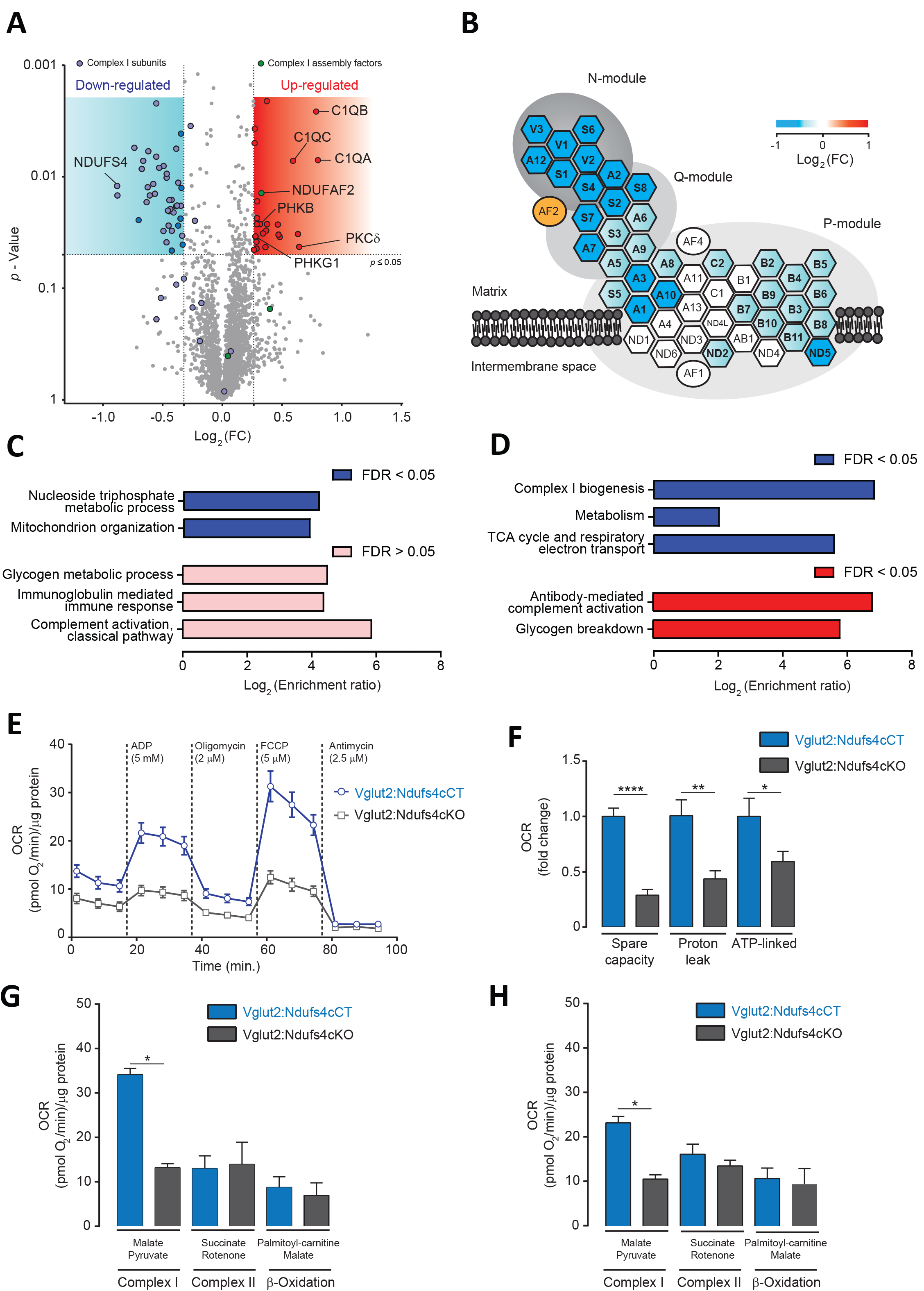
Analysis of the mitochondrial proteome and respiration in mitochondria from the VN of Vglut2:Ndufs4cKO mice. (A) Volcano plot showing proteomic changes induced by NDUFS4-deficiency in mitoTag preparations from Vglut2:Ndufs4cKO and control mice. Blue area contains proteins (n=40) with decreased abundance (<0.8-fold, p≤ 0.05) and red area contains proteins (n=25) with increased abundance (>1.2-fold, p≤ 0.05). (B) Schematic illustration of mitochondrial complex I structure. Heat map shows the changes induced by NDUFS4-deficiency in complex I proteins from Vglut2:Ndufs4cKO mice. Hexagons represent complex I subunits and ovals represent complex I assembly factors. For clarity, many subunits names were shortened by omitting the leading “NDUF”. (C) Enriched biological process GO terms for proteins with decreased (blue) or increased (pink) abundance in mitoTag preparations from the VN of Vglut2:Ndufs4cKO mice. FDR: false discovery rate. (D) Enriched pathways (Reactome) for proteins with decreased (blue) or increased (red) abundance in mitoTag preparations from the VN of Vglut2:Ndufs4cKO mice. (E) Representative pharmacological profile of oxygen consumption rate (OCR) in mitochondria from the VN of Vglut2:Ndufs4cCT and Vglut2:Ndufs4cKO mice. Seahorse assay was run with the assay media containing 10 mM pyruvate and 5 mM malate. OCR was normalized to the total protein amount. (F) Respiratory capacities for mitochondria from the VN of Vglut2:Ndufs4cCT and Vglut2:Ndufs4cKO mice (calculated from Fig.2E). Data are presented as the mean ± SEM. Statistical analysis was performed using an unpaired t-test (\**P* < 0.05, \*\**P* < 0.01, \*\*\*\**P* < 0.0001). (n=10-13). (G-H) OCR of VN mitochondria was measured in the presence of pyruvate/malate, succinate/rotenone, palmitoyl-carnitine/malate, following the addition of ADP, oligomycin, FCCP, and antimycin. Representative OCR results obtained with the addition of FCCP (G) and ADP (H) are shown. Data are presented as the mean ± SEM. Statistical analysis was performed using an unpaired t-test (\*\*\*\**P* < 0.0001).

To explore the functional relevance of the proteomic alterations observed in Vglut2:Ndufs4cKO mice, we conducted bioenergetics analysis on mitochondria preparations from the VN. Mitochondrial oxygen consumption rate (OCR) was assessed by using a Seahorse XF24 extracellular flux analyzer (Fig. 3E). Mitochondria were first incubated with pyruvate and malate as respiratory complex I substrates (state 2). Then, ADP was injected to induce phosphorylation (state 3), followed by oligomycin to inhibit the ATP synthase (state 4) and the chemical uncoupler FCCP to induce maximal respiration (state 3u). VN mitochondria from Vglut2:Ndufs4cKO mice exhibited a profound decrease in the OCR (state 2, 3 and 3u) compared to Vglut2:Ndufs4cCT mice. A more detailed analysis of VN mitochondria respiration (Fig. 3F) revealed significantly reduced spare capacity, proton leak and ATP-linked respiration in Vglut2:Ndufs4cKO compared to control mice. Finally, whether the decrease VN mitochondria bioenergetics was complex-I mediated was investigated. To that end, we used saturating concentrations of specific substrates for complex I (pyruvate/malate), complex II (succinate/rotenone) and ß-oxidation (palmitoyl-carnitine/malate) (Fig. 3F-G). OCR results revealed that state 3 (ADP-induced) and state 3u (FCCP-induced) respirations were similar between Vglut2:Ndufs4cKO and control mice for complex II and ß-oxidation substrates. Collectively, these data show a selective complex I-mediated dysfunction in mitochondria from the VN of Vglut2:Ndufs4cKO mice.

### Hyperacetylation of PDH subunits in VN neurons of Vglut2:Ndufs4cKO mice

Mitochondrial complex I deficiency has been shown to lead to protein hyperacetylation in the heart (Karamanlidis et al., 2013). Hence, we next sought to assess changes in protein acetylation due to NDUFS4 deficiency in mitochondria from glutamatergic neurons in the VN. Acetylated proteins and their modification sites were identified after peptide labeling and affinity enrichment of lysine-acetylated (Kac) peptides by high-resolution LC-MSn in mitoTag preparations from Vglut2:Ndufs4cKO and Vglut2:Ndufs4cCT mice (Figure 4A). A total of 3,445 spectra corresponding to 2,364 nonredundant peptides (928 proteins) were identified through database search (1% FDR). From these, 798 lysine acetylated peptides in 368 protein groups were accurately quantified. A total of 94 differentially acetylated peptides were identified in mitoTag preparations from the VN of Vglut2:Ndufs4cKO mice when compared to controls (p≤ 0.05). Functional protein association network analysis of differentially acetylated proteins using STRING (Fig. 4B) showed that these fit into different GO biological process nodes, being oxidation-reduction process, ATP metabolic process, mitochondrial respiratory chain complex I assembly and pyruvate metabolic process the most significantly enriched biological process categories. Among these proteins, 55 peptides displayed decreased lysine acetylation, and many (n=20) matched to complex I subunits, including NDUFA2, NDUFA5, NDUFA9, NDUFA10, NDUFA12, NDUFAB1, NDUFB3, NDUFB4, NDUFB5, NDUFB6, NDUFB9, NDUFS1 and NDUFV1. Normalization to the total protein abundance confirmed these data. On the other hand, 39 peptides showed increased lysine acetylation, 7 of them corresponding to PDHA1, a subunit that plays a key role in the function of the pyruvate dehydrogenase (PDH) complex. LC-MSn analysis identified 9 sites of lysine acetylation out of all 23 lysine residues on the PDHA1 enzyme (Figure 4C-D). An annotated spectrum of regulated sites of acetylation is provided in Fig. 4C. Among them, 5 sites (K63, K77, K83, K321 and K385) showed increased acetylation in NDUFS4-deficient mitochondria (fold-change> 1.2; p≤ 0.05) (Fig. 4E). To correlate this increased acetylation in specific residues of PDHA1 with changes in the activity of the enzyme, we next assessed mitochondrial preparations from Vglut2:Ndufs4cKO and control mice for PDH activity. Mitochondrial preparations from the VN of Vglut2:Ndufs4cKO mice showed reduced PDH activity when compared to controls (Fig. 4F). No significant changes were observed in the abundance of PDH subunits (PDHA1, PDHB, DLAT, DLD, PDHX) in mitoTag-isolated mitochondria from the VN of Vglut2:Ndufs4cKO mice when compared to controls (Fig. 4G).

**Figure 4.**
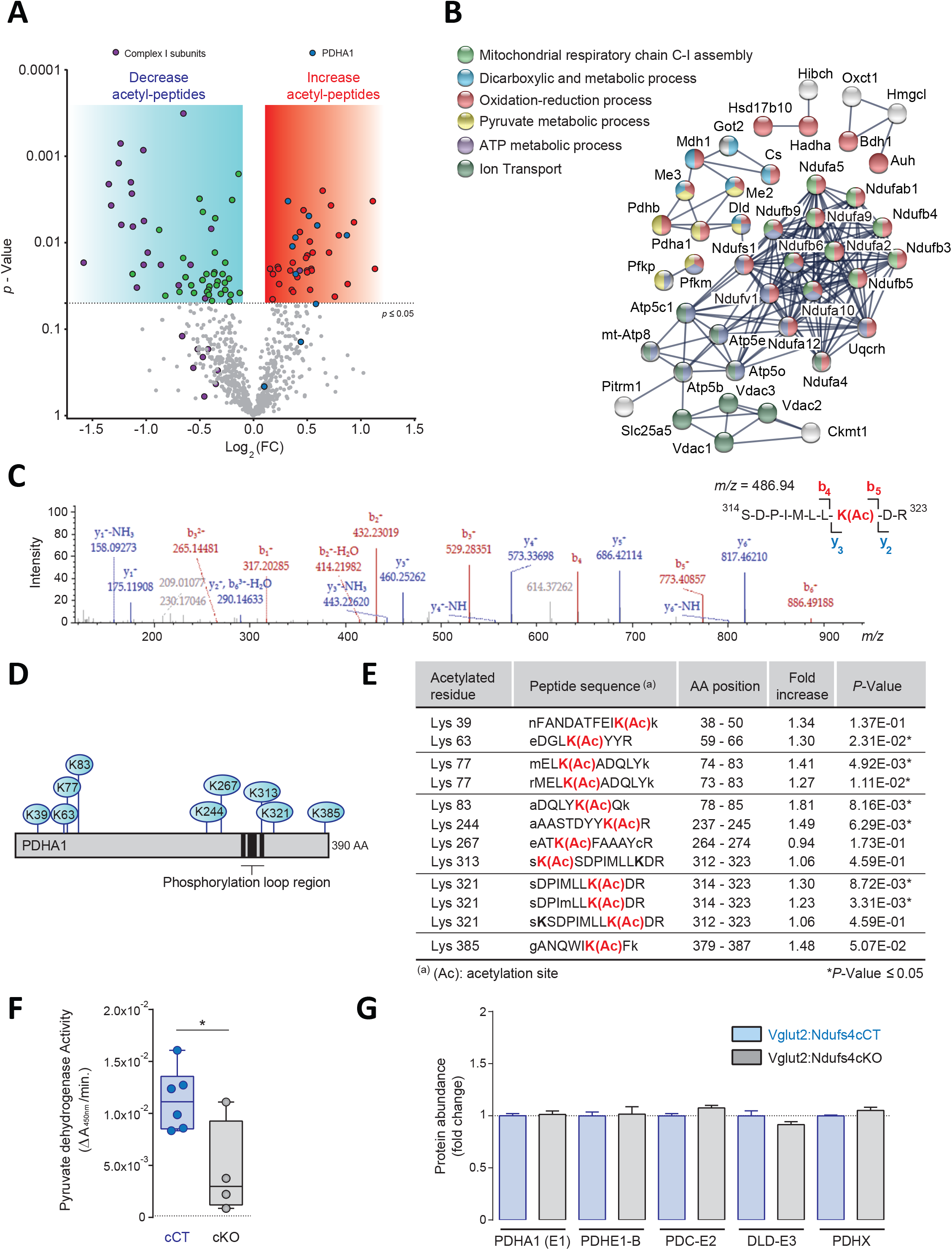
Analysis of the acetylome in NDUFS4-deficient mitochondria from the VN of Vglut2:Ndufs4cCT mice. (A) Volcano plot showing acetyl-peptides with decreased (Blue area; n= 55, p≤ 0.05) or increased abundance (red area; n=39, p≤ 0.05) in mitoTag preparations from the VN of Vglut2:Ndufs4cKO mice. (B) Protein-protein interaction network of proteins with differential acetylation (n= 37) in mitoTag preparations from the VN of Vglut2:Ndufs4cKO mice using STRING. (C) LC-MSn spectrum of the acetylated PDHA1 lysine 321 (in red) containing peptide ^314^SDPIMLLKacDR^323^ (m/z = 486.94), with a mass of 1458,82 Da (D) Schematic representation of the pyruvate dehydrogenase E1 component subunit alfa (PDHA1) showing acetylation sites. (E) Identification of acetylated PDHA1 peptides by mass spectrometry. (E) Pyruvate dehydrogenase activity in mitochondria isolated from the VN of Vglut2:Ndufs4cCT and Vglut2:Ndufs4cKO mice. (G) Relative protein abundance of PDH complex subunits in mitoTag preparations from the VN of Vglut2:Ndufs4cCT and Vglut2:Ndufs4cKO mice.

## 4 Discussion

The mechanisms conferring neuronal vulnerability to MD are currently unknown. However, these have been proven to be highly cell type-specific. Here, we have developed a tool that allows rapid, viral vector-based isolation of mitochondria with cell type specificity and in a spatiotemporally restricted fashion, with the overarching goal of identifying the underlying cellular mechanisms driving neuronal susceptibility to mitochondrial dysfunction. Recently, the urgent need to develop approaches to assess mitochondrial (dys)function in susceptible cell types has led to the generation of different genetic tools to isolate mitochondria with cell-type specificity (Bayraktar et al., 2019; Fecher et al., 2019). These are based on mouse lines that conditionally express, in a Cre recombinase-dependent manner, a tag (GFP or 3XHA-EGFP) that is targeted to the outer mitochondrial membrane to enable immunocapture of mitochondria from specific cell types. However, these approaches are limited by the possibility of unexpected transient expression of Cre recombinase during development in defined cell types (Song and Palmiter, 2018). A significant number of Cre-driver cell lines rely on temporally dynamic promoters that may be transiently active during development, leading to off-target recombination events that compromise the cell type specificity and significantly limiting the application of these tools (Lam et al., 2011; Sanz et al., 2015). Cre-dependent viral vector-based approaches such as our mitoTag assay allow delivery of the transgene into the adult brain, therefore overcoming this important limitation. In addition, viral vector-based approaches avoid complex and time-consuming breeding strategies and enable temporal and regional control, therefore allowing rapid and anatomically restricted mitoTag assays.

Our proteomics data shows that NDUFS4 deficiency leads to mitochondria depleted of most of the subunits of the respiratory chain NADH dehydrogenase (complex I; CI). Complex I (CI; NADH: ubiquinone oxidoreductase; EC 1.6.5.3) is essential for oxidative energy metabolism as it contributes to the generation of the proton motive force needed for ATP synthesis (Hirst, 2013). This complex has a boot-shaped structure with a hydrophilic matrix arm and a hydrophobic membrane arm that is embedded in the inner mitochondrial membrane. These arms are subdivided into three structurally and functionally defined modules: the N-module (involved in NADH binding and oxidation; where NDUFS4 is located), the Q-module (for transfer of electrons along Fe-S clusters to ubiquinone), and the P-module (mediating proton pumping)(Formosa et al., 2018). Our results evidence that components of the N-module were the most significantly downregulated in mitochondria from the VN of Vglut2:Ndufs4 mice, indicating the critical role of this accessory subunit is critical for the assembly and/or stability of the N module component of CI, confirming previous studies in LS patients harboring NDUFS4 mutations (Scacco et al., 2003) and *Ndufs4-deficient* mice (Calvaruso et al., 2012; Leong et al., 2012; Valsecchi et al., 2012). Furthermore, our data also shows an increase abundance of a specific assembly factor, NDUFAF2 (B17.2L), a molecular chaperone associated with a large subassembly of ~830-kDa form of complex I and required in the late stage of CI assembly that is essential for the efficient assembly of complex I (Ogilvie et al., 2005; Lazarou et al., 2007; Sanchez-Caballero et al., 2016). This increase may result from a compensatory attempt to keep complex I assembled upon NDUFS4 deficiency.

Our analysis of mitochondria also revealed mitochondrial association of characterized non-mitochondrial proteins upon NDUFS4 deficiency, which would remain unnoticed if a global analysis of the cellular proteome is performed. Vglut2:Ndufs4cKO mitochondria showed increased abundance of the three different polypeptide chains that compose the complement subcomponent C1q. C1q has been shown to be present within the cell (Ten et al., 2010; Ghebrehiwet et al., 2012), and one of the receptors for C1q, the globular head C1q receptor (gC1qR), is a mitochondrial cell-surface protein known to modulate metabolism through association with C1q (Dedio et al., 1998; Kolev and Kemper, 2017). In this regard, an effect on spare respiratory capacity and oxygen consumption has been shown by gC1qR-bound C1q, which is known to enhance nuclear transcription of mitochondrial biogenesis genes (Ling et al., 2018). Association of C1q with NDUFS4-deficient mitochondria may be a mechanism of the cell to promote mitochondrial biogenesis to overcome mitochondrial dysfunction. Conversely, C1q has also been shown to mediate mitochondrial ROS production by cortical neurons in neonatal hypoxic-ischemic brain injury and that C1q-deficient brain mitochondria present preserved activity of the respiratory chain (Ten et al., 2010), suggesting tissue- and likely cell type-specific mitochondrial roles of complement factors. Therefore, the role of C1q association to NDUFS4-deficient mitochondria warrants further research.

Furthermore, our direct proteomics approach has detected increased abundance of PKCδ in mitoTag preparations from Vglut2:Ndufs4cKO mice, suggesting mitochondrial translocation of this kinase in NDUFS4-deficient vulnerable neuronal populations. In this regard, PKCδ has been shown to translocate to mitochondria upon different stressors and in different neurodegenerative pathologies, including oxidative (Majumder et al., 2001) and genotoxic stress (Lasfer et al., 2006), or cerebral ischemia (Dave et al., 2011). Targeting of PKCδ to mitochondria alters the mitochondrial membrane potential and induces apoptotic responses of cells to oxidative stress. Consistently, treatment with pan-PKC inhibitors increases survival and delays the onset of neurological symptoms in NDUFS4KO mice (Martin-Perez et al., 2019). Administration of isoform-specific PKCδ inhibitors to Ndufs4-deficient mice will define the relevance of this protein as a target for the treatment of mitochondrial disease.

CI-deficiency in the heart of conditional NDUFS4KO mice results in a NAD(H) redox imbalance, with a decreased NAD^+^/NADH ratio, inhibiting the activity of the mitochondrial sirtuin deacetylase SIRT3, and leading to protein hyperacetylation (Karamanlidis et al., 2013). However, the effect of NDUFS4-deficiency on the mitochondrial acetylome of neurons susceptible to MD was unknown. Analysis of the mitochondrial acetylome in mitoTag preparations greatly reduces the complexity of the sample by avoiding the interference of heavily acetylated cytosolic and nuclear proteins. This results in increased identification of acetylated peptides corresponding to mitochondrial proteins. We report increased abundance of acetylated peptides corresponding to PDHA1 in NDUFS4-deficient mitochondria. PDHA1 is the E1 alpha 1 subunit containing the E1 active site of the pyruvate dehydrogenase (PDH) complex, involved in transforming pyruvate into acetyl-CoA that is subsequently used by both the citric acid cycle and oxidative phosphorylation to generate ATP. Increased acetylation at specific PDHA1 sites anticipated reduced activity of the enzyme since acetylation is typically reported to inhibit the catalytic activity of mitochondrial enzymes (Ghanta et al., 2013). More specifically, it has been described that acetylation of PDHA1 at K83 and K321 reduce the activity of the enzyme. Noteworthy, both K83 and K321 residues are SIRT3 substrates (Fan et al., 2014; Wei et al., 2019). Therefore, reduced PDH activity may be contributing to the Warburg effect by enhancing glycolysis over mitochondrial oxidation of pyruvate (Warburg, 1956; Kroemer and Pouyssegur, 2008; Vander Heiden et al., 2009). Hence, the reduction in PDH activity may be involved in the increased glycolysis observed in Ndufs4KO mice (Johnson et al., 2013) and contribute to the metabolic demise of *Ndufs4-deficient* cells. Noteworthy, mutations in the PDH gene are associated with X-linked Leigh syndrome (Ruhoy and Saneto, 2014), indicating its central role in mitochondrial function.

To date, the marked heterogeneity in mitochondrial composition and function has limited our understanding of the underlying molecular determinants of the neuronal susceptibility to mitochondrial dysfunction. Therefore, it is paramount to study mitochondrial function in a genetically-defined manner to unravel the pathogenic mechanisms of mitochondrial disease. Here, by combining mouse genetics and cell type-specific mitochondrial isolation, we have assessed the mitochondrial proteome of a neuronal population shown to be highly susceptible to mitochondrial dysfunction in a well-established mouse model of mitochondrial disease. Our results underscore the marked consequences of CI-deficiency in the proteome of affected neurons, identifying the metabolic consequences of such alterations. Furthermore, we have identified a potential role for PDH, the complement factor C1q, and PKCδ in the pathology, paving the way for future studies of the therapeutic potential of modulating these pathways in the quest for effective treatments of mitochondrial disease.

## 6 Author Contributions

AG, PPD, MC, MGT, JA, ES, AQ conducted experiments and acquired and analyzed the data. AG, MC, JA, ES and AQ designed research studies. AG, ES, AQ wrote the manuscript. AQ and JA provided reagents and ES and AQ coordinated the work.

## 8 Acknowledgments

We are also indebted to Dr. Alejandro Sànchez and Ms. Núria Barba from ‘‘Servei de Microscopia” from Universitat Autònoma de Barcelona (UAB) for helpful technical assistance (http://sct.uab.cat/microscopia)

